# Poor decision making and sociability impairment following central serotonin reduction in inducible TPH2-knockdown rats

**DOI:** 10.1101/2024.01.06.574479

**Authors:** Lucille Alonso, Polina Peeva, Tania Fernández del valle Alquicira, Narda Erdelyi, Ángel Gil-Nolskog, Michael Bader, York Winter, Natalia Alenina, Marion Rivalan

**Affiliations:** Humboldt-Universität zu Berlin, Berlin, Germany; Charité – Universitätsmedizin Berlin, corporate member of Freie Universität Berlin and Humboldt-Universität zu Berlin, Berlin, Germany; Univ. Bordeaux, CNRS, IINS, UMR 5297, F-33000 Bordeaux, France; Max Delbrück Center for Molecular Medicine in the Helmholtz Association, Berlin, Germany; DZHK (German Center for Cardiovascular Research), Partner Site Berlin, Berlin, Germany; Institute for Biology, University of Lübeck, Lübeck, Germany; NeuroPSI – Paris-Saclay Institute of Neuroscience, CNRS - Université Paris-Saclay, Saclay, France

**Keywords:** TPH2, serotonin, tetracycline responsive system, inducible knock-down, vulnerability, network analysis, rat

## Abstract

Serotonin is an essential neuromodulator for mental health and animals’ socio-cognitive abilities. However, we previously found that a constitutive depletion of central serotonin did not impair rat cognitive abilities in stand-alone tests. Here we investigated how a mild and acute decrease of brain serotonin would affect rats’ cognitive abilities. Using a novel rat model of inducible serotonin depletion via the genetic knock-down of tryptophan hydroxylase 2 (TPH2), we achieved 20% decrease of serotonin levels in the hypothalamus after three weeks of non-invasive oral doxycycline administration. Decision making in the Rat Gambling Task and the Probability Discounting Task, cognitive flexibility and social recognition memory were tested in low-serotonin (Tph2-kd) and control rats. Our results showed that the Tph2-kd rats were more prone to choose disadvantageously on the long term (poor decision making) and that only the low-serotonin poor decision makers were more sensitive to probabilistic discounting and had poorer social recognition memory than other low-serotonin and control individuals. Flexibility was unaffected by the acute brain serotonin reduction. Poor social recognition memory was the most central characteristic of the behavioral network of low-serotonin poor decision makers, suggesting a key role of social recognition in expression of their profile. The acute decrease of brain serotonin appeared to specifically amplify cognitive impairments of the subgroup of individuals also identified as poor decision makers in the population. This study highlights the great opportunity Tph2-kd rat model offers to study inter-individual susceptibilities to develop cognitive impairment following mild variations of brain serotonin in otherwise healthy individuals. These transgenic and differential approaches together could be critical for the identification of translational markers and vulnerabilities in the development of mental disorders.

## Introduction

Mental health is a dynamic process and alternation between phases of deterioration and improvement of health is one hallmark of mental illness. In regards to the current lack of specific and universal treatments of mental disorders (Moriana et al., 2017), identifying individual-specific targets to prevent transition from adaptive to pathological mental states, is essential (World Health Organization, 2021). Along the continuum between adaptive and maladaptive behavioral dimensions, vulnerable individuals at higher risk for the development of psychiatric disorders, would present a combination of interconnected preserved and impaired behaviors (Dalgleish et al., 2020). Following the network approach to mental disorder a higher connectivity within such networks of symptoms is an objective characteristic of vulnerability to pathology (Borsboom, 2017). In humans, making repeated poor decisions in the everyday life is known to lead to long-term disadvantageous outcomes and a general deterioration of mental health. Poor decision making abilities is one common symptom to most human mental disorders (Cáceda et al., 2014). In the Iowa Gambling Task, a test for decision making in everyday uncertain conditions of choices, around 30% of non-clinical populations present similar decision making deficits as clinical populations (Bechara and Damasio, 2002; Denburg et al., 2005; Steingroever et al., 2013). In previous studies, we have identified a subpopulation of healthy rats whose primary deficit consist in less advantageous strategies of choice in uncertain and complex conditions of choice as tested in the Rat Gambling Task (Alonso et al., 2019; Rivalan et al., 2009). Healthy poor decision makers are consistently found, between labs (Daniel et al., 2017; Mu et al., 2015), across strains (WH and DA (Alonso et al., 2019), SD (this paper and (Daniel et al., 2017; Mu et al., 2015) and Long Evans –unpublished data–) and species (mouse (Pittaras et al., 2016), primate and monkey (Proctor et al., 2014)). Healthy rats’ poor decision makers present a unique combination of preserved and impaired neurological and behavioral characteristics (Alonso et al., 2023, 2019; Rivalan et al., 2013, 2011, 2009). They express high reward-seeking and risk-seeking behavior, inflexibility, and a tendency to dominant behavior together with normal control of cognitive impulsivity and economical risk taking and a normal social everyday life (Alonso et al., 2019; Rivalan et al., 2013). In addition, poor decision makers exhibit an imbalance in brain monoaminergic neurotransmitters and a smaller and weaker cortical-subcortical brain network activated during the Rat Gambling Task (Fitoussi et al., 2014). With such a vulnerable profile it is however not known if poor decision makers are indeed more vulnerable to an acute biological change in the way that it would impair their phenotype more than the phenotype of other individuals.

Central serotonin is an essential neuromodulator for mental health and a promising transdiagnostic marker of mental illness. Selective serotonin reuptake inhibitors can be prescribed following adverse life events in order to boost the central serotonergic system and improve coping processes facing for example grief, unsolicited work-termination, or seasonal affective disorder (Bui et al., 2012; Hyde et al., 2015; Nussbaumer-Streit et al., 2021). In animals, the models of choice to cause an acute biological perturbation are genetic On-Off inducible models (Kotnik et al., 2009). The new TetO-shTPH2 transgenic rat model (Matthes et al., 2019; Sidorova et al., 2021) is a knockdown model that targets central serotonin to create a mild acute drop of central serotonin. The application of Doxycycline (Dox) in the drinking water of TetO-shTPH2 rats induces the expression of shRNAs against messenger RNA of tryptophan hydroxylase 2 (TPH2) which results in a decrease up to 25% of brain serotonin levels (Matthes et al., 2019). In the current study, we used TetO-shTPH2 rats to test the impact of acute moderate brain serotonin imbalance on cognition and depending on individual spontaneous decision making type. We focused on complex decision-making and risky based decision making, cognitive flexibility, and social recognition memory which are serotonin-dependent transdiagnostic symptoms of psychiatric disorders. We explored how these functions interacted with each other using a novel behavioral network analysis. Following our hypothesis of increased vulnerability in poor decision makers, we expected the moderate drop of serotonin function to impair more specifically their behavior and that it would also reflect on the properties of their behavioral network.

## Material and methods

### Ethics

All procedures followed the national regulations in accordance with the European Union Directive 2010/63/EU. The protocols were approved by the local animal care and use committee (LaGeSo, Berlin) and under the supervision of the animal welfare officer of our institution.

### Animals

We used 96 male TetO-shTPH2 transgenic rats of Sprague Dawley background (SD) and 24 male SD rats, Control animals, included 36 TetO-shTPH2 rats treated with water (TetO-water) and 24 SD rats treated with Dox (SD-Dox). Tph2-knockdown animals, called Tph2-kd later on, included the 60 TetO-shTPH2 rats treated with Dox (TetO-Dox). Number of animals for each test and exclusion criteria are reported in Figure 1. Animals were tested by group of 12 individuals called a batch and consisting of 6 Controls and 6 Tph2-kd. Eight out of ten batches of animals were tested at the same time in pairs. Batches are indicated in Figure 1 and pairs of batches in Figure S1.

**Figure 1.**
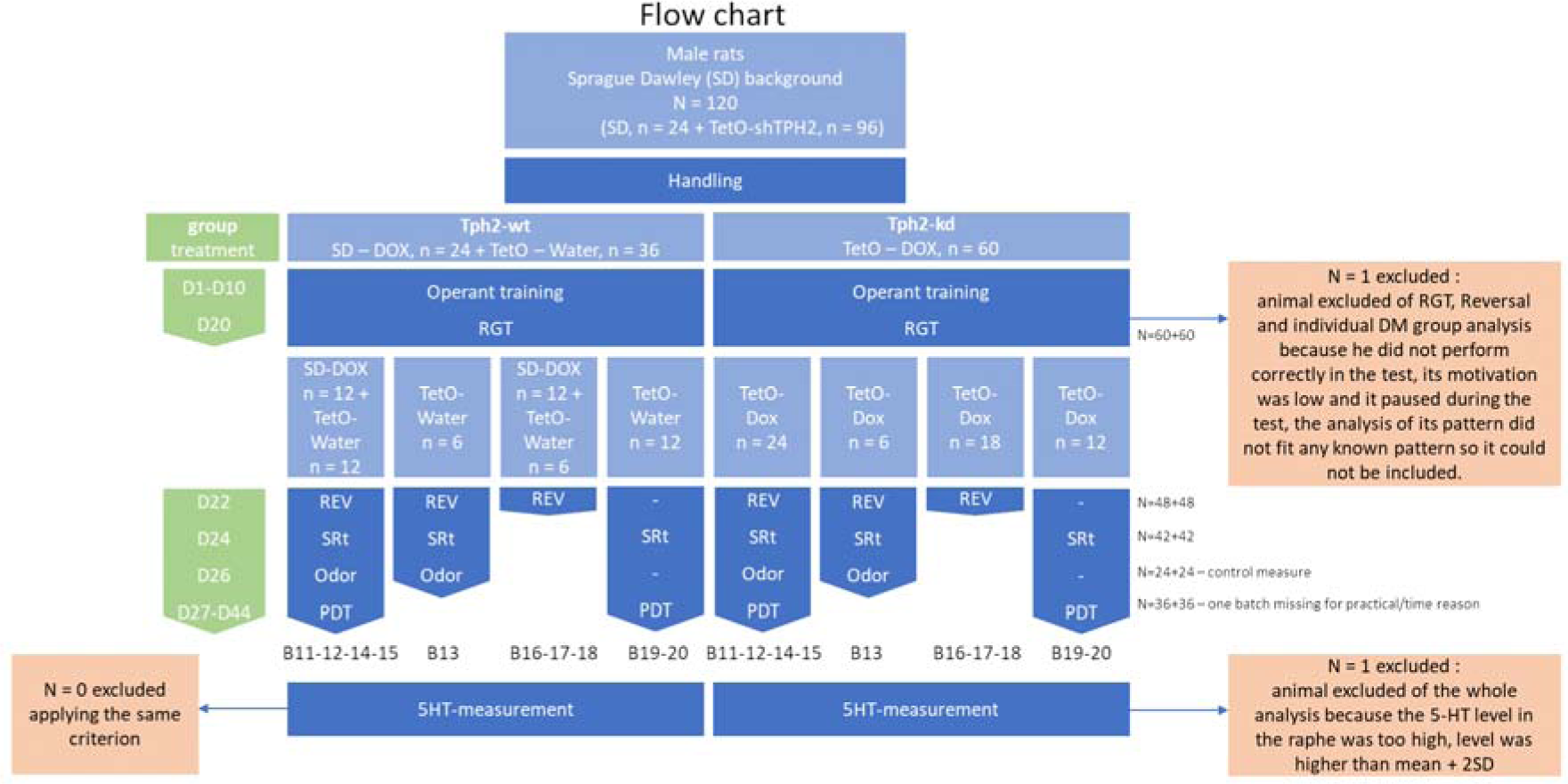
Flow chart for reporting attrition.

Animals were born at the Max Delbrück Center for Molecular Medicine, Berlin and transferred to the animal facility of Charité – Universitätsmedizin Berlin between nine and ten weeks of age. They were housed in standard rat cages (Eurostandard Type IV, 38 cm x 59 cm) in pairs of animals receiving the same treatment (TetO-water, SD-Dox or TetO-Dox). Cages were maintained in temperature-controlled rooms (22°C–24°C and 45%–55% humidity) with inverted 12-hour light-dark cycles. Animals had *ad libitum* access to water and to standard maintenance food (V1534-000, Ssniff, Germany). During operant training and testing, they were maintained at 95% of their free-feeding weight. After their daily operant testing rats were fed up to 40 g per animal depending on the amount of reward (sweet pellets) they received in the operant chamber and following an unpredictable schedule (one to several hours after the end of test) to avoid their anticipation of feeding. Rats were weighed every two to three days allowing for adjustment of their portion of standard food and drug treatment.

### Treatment

Animals received Dox from the first day of operant training and until the end of the protocol in the drinking water of the cage (Fig.1). The Dox solution was prepared every three days at a dosage of 40 mg/kg of body weight. The concentration of the solution was adapted to the average water consumption and body weight of the animals. For the batches tested in pairs, Dox solutions were prepared from the same supply bottle by the same experimenter and at the same time.

### Sacrifice and brain collection

Two days after the last test, whatever the length of the protocol, rats were anesthetized via an intraperitoneal injection of Ketamine (100mg/kg) and Xylazine (10mg/kg) under isoflurane anesthesia. Pairs of batches were sacrificed by the same experimenters over two days. Animals were transcardially perfused with phosphate-buffered saline. Brain parts were immediately collected, snap-frozen on dry ice, and stored at −80 °C until further use. Orbitofrontal area, prefrontal area, hippocampus, hypothalamus, and raphe were dissected. Brain tissue was weighted after freezing. After the first pilot batches tested in pair (11&12), the time of the sacrifice was controlled for to prevent from circadian effects on the measurements. Sacrifice started one hour after the start of the dark phase except for batch 12 for which it started five hours before the dark phase.

### HPLC analysis

For the determination of the dosages of monoamines and their metabolites in brain tissue, frozen tissues were homogenized in 300 µl lysis buffer containing 10 µM ascorbic acid and 1.8 % perchloric acid using a FastPrep system. Samples were centrifuged for 30 min at 13,000 rpm. Supernatants were transferred in Eppendorf tubes and stored at -80 °C until HPLC measurement. Tissue levels of TRP, 5-HT and its metabolite 5-HIAA were analyzed using high sensitive HPLC with fluorometric detection (Shimadzu, Tokyo, Japan). Sample separation took place at 20□°C on a C18 reversed-phase column (OTU LipoMareC18, AppliChrom Application & Chromatography, Oranienburg, Germany) using a 10□mM potassium phosphate buffer, pH 5.0, containing 5% methanol with a flow rate of 2□ml□min–1.

Calculation of substance levels was based on external standard values. Amounts of 5-HT, 5-HIAA, and TRP were measured in hypothalamic samples and normalized to the wet tissue weight for statistical analysis. Individual concentrations of 5-HT, 5-HIAA and TRP were normalized per batch to the mean of the control individuals and are presented as percentage.

### Behavioral testing

Animals were grouped in batches of 12 animals (6 Tph2-wt and 6 Tph2-kd) and were tested either in the morning or in the afternoon (i.e. 24 animals per day) depending on the light cycle of the housing room (lights on at 20:00 in room 1 or 01:00 in room 2) in order to maximize the use of our four operant cages and minimize potential circadian effect. Rats were all tested 1h after start of dark phase and within less than 3h (5h for the social recognition test).

### Operant system

The four operant cages (Imetronic, France) contained on one side a curved wall equipped with two or four nose-poke holes, depending on the test. On the opposite wall was a food magazine connected to an outside pellet dispenser filled with 45 mg sweet pellets (5TUL Cat#1811155, TestDiet, USA). A separator with a 10 x 10 cm aperture was placed in the middle of the cage. The same light conditions applied to the four cages.

### Rat gambling task (RGT)

For the RGT, operant cages were equipped with four nose-poke holes arranged in two pairs (5 cm between holes) on each sides of the curved wall (12.5 cm between pairs). The training and testing procedures (Alonso et al., 2023, 2019; Rivalan et al., 2013, 2009) were adapted to the treatment period required by the tetO system. The training started at day 1 of the Dox treatment (Fig.1). First, rats learned to poke into the nose-poke holes and retrieve the associated reward (1 pellet) into the magazine. Training 1 was completed when 100 pellets had been collected in 30 min (cut-off). Training 2 consisted in poking two consecutive times into the same hole to get a reward (1 pellet). Training 2 was completed when 100 pellets had been collected in 30 min (cut-off). Two consecutive nose-pokes into the same hole were considered as a choice for all operant tests. Then, training 3a consisted of a short session during which a choice to any hole was rewarded with 2 pellets within 15 min (cut-off and 30 pellets maximum). After that, a forced training (Alonso et al., 2019) was given to those rats who had a preference above 60% for one of the two pairs of holes in the last session of training 2. During the first part of the forced-training, the two nose-poke holes on the non-preferred side were active and lit; the two holes on the preferred side were inactive and not lit. Choosing the active holes induced the delivery of one pellet. After the collection of 15 pellets, the second part of the forced training started with the four holes active and lit. Choosing holes of the formerly preferred side induced the delivery of one pellet with a probability of 20% whereas choosing the formerly non-preferred side induced the delivery of one pellet with a probability of 80%. The cut-off was 50 pellets or 30 minutes. The whole training was completed in seven to ten days. Rats were fed *ad libitum* from training completion until day 18 of the treatment (Fig.1). On day 19 a session of training 2 was done in order to check the behavior of the rats and any side preference. A forced training was applied on the same day to the rats with a preference for one side superior to 80% (n = 4 animals). This criterion was refined depending on which side the preference was developed for the last three cohorts, the preference threshold was 80% for the side that will be advantageous during the test and 70% for the side that will be disadvantageous (n = 3 animals).

Testing took place on the twentieth day of treatment for 60 minutes. Two nose-poke holes on one side were rewarded with a large reward (two pellets) and associated with unpredictable long time-outs (222s and 444s with the probability of occurrence ½ and ¼ respectively). This was the disadvantageous option, leading to a lower maximum gain of pellets in 60 minutes. Two nose-poke holes on the other side were rewarded by one pellet and associated with unpredictable short time-outs (6s and 12s with the probability of occurrence ½ and ¼ respectively). This was the advantageous option, leading to a maximal gain of pellets within 60 minutes. The percentage of advantageous choices for the last 20 minutes of RGT was used to identify the sub-types of decision makers: good decision makers (GDMs) with more than 70% of advantageous choices, poor decision makers (PDMs) with less than 30% of advantageous choices and intermediate animals in between. Computed over the 60 minutes of testing, the percentage of advantageous choices per ten-minute interval indicated the progression of the preference over time. An index of the motivation for the reward was measured as the mean latency to visit the magazine after a choice.

### Reversed-RGT

The animals were tested in the reversed-RGT 48 hours after the RGT. For this test the two disadvantageous options were spatially switched with the two advantageous options (Alonso et al., 2023, 2019; Rivalan et al., 2013). A flexibility score was calculated as the preference during reversed-RGT for the location of the non-preferred option during the RGT. Flexible rats had more than 60% of such choices during the last 20 minutes, undecided rats had between 60% and 40% choices, and inflexible rats had less than 40% choices.

### Probability discounting task (PDT)

For the PDT, operant cages were equipped with two nose-poke holes on each outer side of the curved wall (25 cm between holes). The protocol was adapted to reduce its overall duration (Alonso et al., 2023) due to the pharmacological treatment. One hole (NP1) was associated with a small and sure reward (1 pellet), the second one (NP5) was associated with a large and uncertain reward (5 pellets) (Koot et al., 2012). The delivery of the large reward was dependent on the probability applied that changed from high to low during the experiment: P = 1, 0.33, 0.20, 0.14 and 0.09. The probability was fixed for a day and increased the next day only after reaching the stability criterion. At least three training sessions (P = 1) were done and a percentage of choice of the large reward ≥ 70% on two following sessions with ≤ 15% variation was required to start the test. During testing sessions (P < 1) of the stability criterion was ≤ 10% variation of choice of the large reward during two consecutive sessions. The percentage of preference for the large and uncertain reward was calculated for each probability as the percentage of NP5 choices during the two stable sessions. To calculate the area under the curve (AUC) which represents the sensitivity to probabilistic uncertainty and risk taking, for each individual the preference for the large reward for each probability was normalized to the preference for the large reward during the training (P = 1) and plotted versus the probabilities expressed as odds with odds = (1/P) -1 (Zoratto et al., 2014).

### Social recognition task (SRt) (Alonso et al., 2023)

The test took place in a square open field (50 x 50 cm). A small clean and empty cage was placed in one corner of the open field shielded by walls to avoid the test rat hiding behind the cage. The unfamiliar conspecifics were older Wistar Han rats, accustomed to the procedure. A video camera on top of the OF recorded the experiment. The subject was placed in the OF containing the empty cage for a habituation of 15 minutes. Then, the unfamiliar conspecific was placed in the small cage and the subject was allowed to freely explore the open field for five minutes (E1). After that the small cage with the conspecific was removed from the open field, a second clean and empty cage was used to fill the space and the subject remained alone in the open field for a break of 10 minutes. The encounter procedure was repeated two more times with the same conspecific (E2, E3). The time spent in close interaction with the unfamiliar rat was measured for each encounter and for the first five minutes of Habituation (Hab) when the subject smelled at the grid of the empty cage. The social preference was calculated as the ratio of the interaction time in E1 and Hab. The short-term social recognition memory was calculated as the ratio of the interaction time in E1 and E3.

### Odor discrimination test (Alonso et al., 2023)

The test took place in a square OF (50 x 50 cm). Two plastic petri dishes filled with either used or fresh bedding were placed in two opposite corners of the OF. Except for the first group, the dishes were taped onto the OF flour to avoid animals pushing them around. A video camera above the OF recorded the experiment. The test rat explored the OF for 5 min. The time spent in close interaction with each dish was measured by trained observers using JWatcher (version 1.0) (Blumstein et al., 2000) and the preference for the used bedding (social odor) was calculated.

### Statistical analysis

The free software R (version R-3.6.1) and R studio (version 1.1.456) were used for the statistical analyses (R Core Team, 2019). We used the Wilcoxon rank sum test to compare the Tph2-wt and Tph2-kd groups, the Kruskal-Wallis rank sum test to compare the decision maker groups against each other (GDMs, INTs and PDMs of Tph2-wt and Tph2-kd groups), Fisher’s exact test to compare the number of GDMs and PDMs in each groups, and the Wilcoxon sign test (Package RVAideMemoire) (Hervé, 2020) to compare the performance of the animals to a theoretical value in PDT, SRt and odor discrimination test. We used ANOVA with permutations (package lmPerm) (Wheeler and Torchiano, 2016) which are suited for small groups and non-parametric data to compare the multiple groups and decision maker groups (dm-group) over several time points (probability, encounter) with animal as error factor.

### Network visualization of behavioral data

We used a network analysis method (Package qgraph) (Epskamp et al., 2012) to visually represent the strength of connections between pairs of behaviors, as behavioral networks of the groups and subgroups. According to the network approach of psychopathology, the visual representation of the connectivity between symptoms could inform about the potential dynamic existing between them in pathological context. In the current study we used this visualization to explore how the five tested cognitive functions connected to each other depending on their treatment and decision making profile. We also were interested to see if one function would appear more central (in terms of number and weight of ties to other nodes) than the others.

The five parameters constituting the nodes of the networks were the RGT score, mean latency to visit the magazine after a choice during RGT, AUC of the PDT, social preference, and short-term social recognition memory. They represented complex decision making ability, motivation for reward, risky decision making ability, social preference and social recognition memory respectively. The reversed-RGT and odor discrimination tests were not performed by all animals and were not included in the networks. The network analysis was done using the qgraph() function (type of graph “association”) where correlations are used as edge weights between two nodes. Although partial correlations are usually preferred to correlations as they account for the relationships of the network for each pairwise link, it was not possible in our analysis to apply Spearman’s partial correlation to the Tph2-kd poor decision maker group as the number of individuals equals the number of functions. We used the Spearman’s correlations (package Hmisc) for all groups. To simplify the description and visualization of the networks only correlations above 0.28 were represented, and the networks were calibrated from 0 to 0.7. We calculated the strength centrality for each nodes taking which is the centrality of a node taking into account the number and the weight of edges connecting to the nodes (Opsahl et al., 2010). With only one control poor decision maker individual no network could be computed for this subgroup.

## Results

We first checked if Dox treatment effectively decreased serotonin metabolism in Tph2-kd rats. The serotonin level in the hypothalamus was lower in the Tph2-kd group compared to the control group (Fig.2A, Wilcoxon rank sum test with continuity correction, W = 2259, p-value < 0.001). The Dox treatment induced in the Tph2-kd rats a decrease of serotonin levels of 21% in average (sd = 23). The level of 5-HIAA was also decreased by 25% in average (sd = 23, Fig.2B, Wilcoxon rank sum test with continuity correction, W = 2825, p-value < 0.001) whereas as expected the tryptophan level was stable between groups (Fig.2C, Wilcoxon rank sum test with continuity correction, W = 2034, p-value = 0.1141). The ratio of 5-HIAA/TRP indicated a consistent decrease of serotonin metabolism in Tph2-kd animals independent of the duration of the treatment (Fig.S1, p.anova, *treatment*, F(1,107) = 75, p-value < 0.001, *duration*, F(1,7) = 0.18, p-value = 0.719) and showed variation in the 5-HIAA/TRP decrease between batches. Due to a technical problem, for batch 12, 5-HT could not be correctly measured, nevertheless the 5-HIAA/TRP ratio showed the effect of the treatment for this group (Fig.S1).

**Figure 2.**
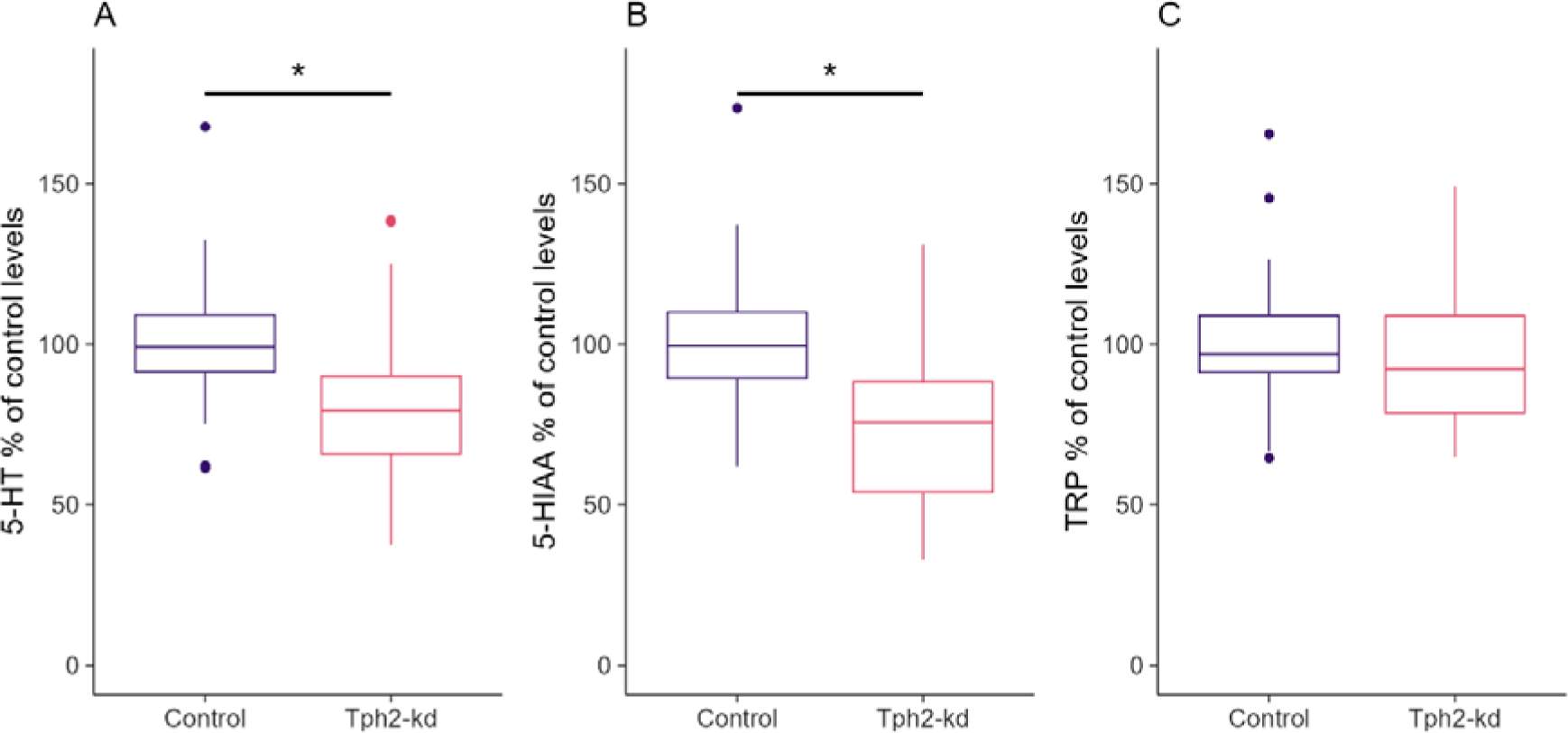
Levels of **A.** Serotonin (5-HT) **B.** 5-HIAA and **C.** Tryptophan levels in the hypothalamus in % of controls. Values were normalized to control individuals within each batch, one batch was excluded from **A** due to a technical problem of serotonin detection. Boxplots classically represent the median, 25th and 75th percentiles and 1.5 interquartile range (IQR). * p-value < 0.05. Panel A: Tph2-wt, n = 54, Tph2-kd, n = 54. Panels B-C: Tph2-wt, n = 60, Tph2-kd, n = 59.

In both groups, each animal showed either one of the three typical decision making strategies (Fig.3A). All animals started the test at 50% of advantageous choices (Fig.3A, Kruskal-Wallis rank sum test, chi-squared = 10.2, df = 5, p-value = 0.067, after 10min: Tph2-wt-gdm 56 + 17 (mean + SD), Tph2-wt-int 52.5 + 17, Tph2-wt-pdm 38.5 + 24, Tph2-kd-gdm 59.6 + 17, Tph2-kd-int 52 + 11, Tph2-kd-pdm 39.2 + 18). For good decision makers (upward triangles) the percentage of advantageous choices increased until reaching a very high preference at the end of the test (above 70%). For poor decision makers (downwards triangles) the percentage of advantageous choices decreased until reaching a very low preference at the end of the test (below 30%). For intermediate individuals (squares) the percentage of advantageous choices stayed around 50% until the end of the test (between 70% and 30%). More disadvantageous choices (RGT score < 30%) were made by Tph2-kd rats than by Tph2-wt rats (Fig.3B-left, Wilcoxon rank sum test, W = 2112.5, p-value = 0.045). The difference between Tph2-wt and Tph2-kd groups was found in the proportion of individuals in each decision making category with a higher proportion of poor decision makers in Tph2-kd than the Tph2-wt group (Fig.3B-right, Fisher’s exact test, p-value = 0.044). The distribution density of each group similarly illustrates the decrease in individuals showing good decision making strategy after acute serotonin drop in Tph2-kd animals (Fig.3B-right). The latency to collect the reward was dependent on the decision making group (Fig.S2, p.anova, *RGT score*, F(1,115) = 38, p-value < 0.001) and independent of the treatment group (Fig.S2, p.anova, *treatment*, F(1,115) = 0.14, p-value < 1.103).

**Figure 3.**
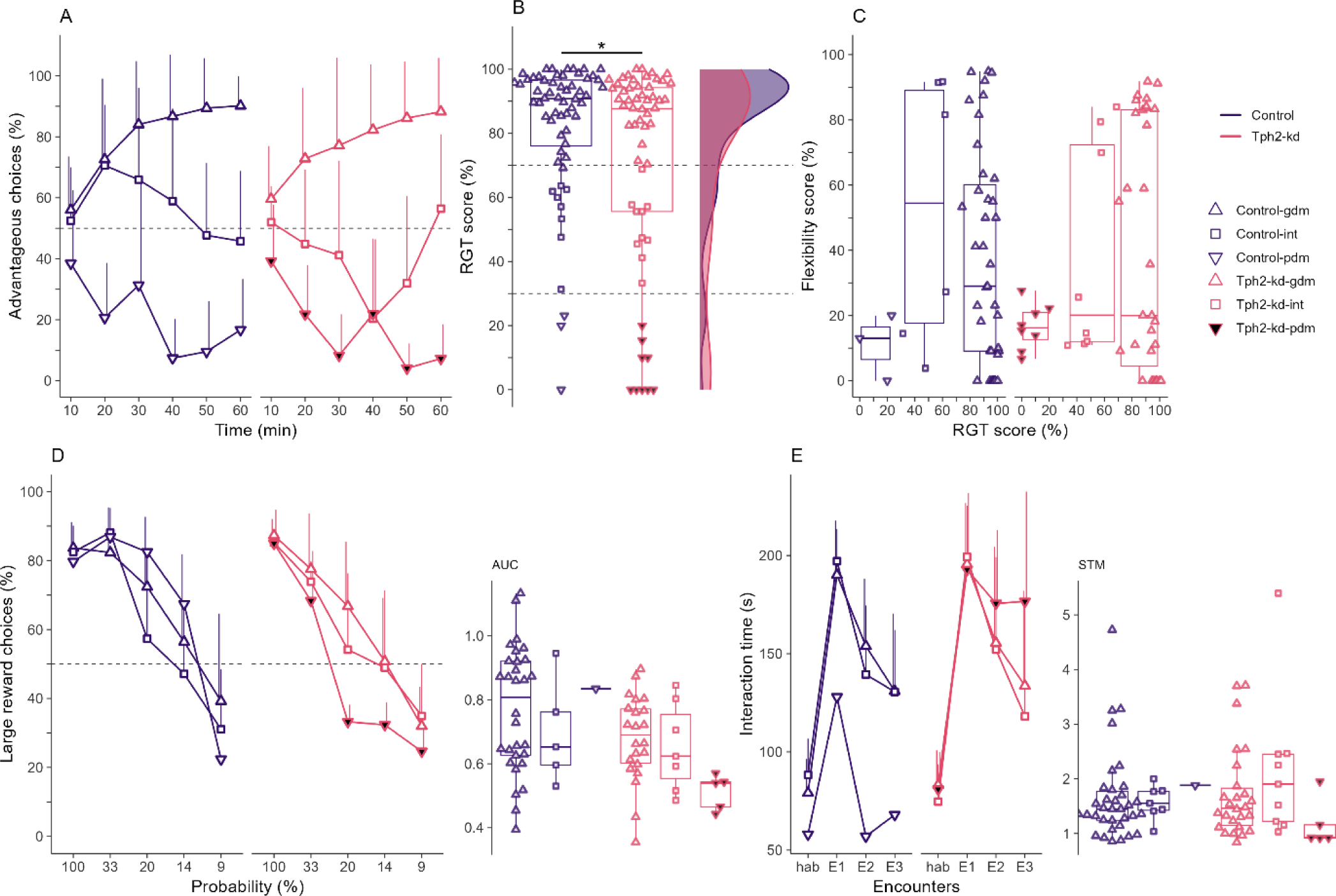
**A.** Advantageous choices in the rat gambling task (RGT) over time (10-minute bins). Lines indicate mean + SD, dashed line shows 50% chance level. **B. Left** Individual (mean) scores during the last 20 minutes of the RGT (RGT score). The dashed lines at 70% and 30% of advantageous choices visually separate good decision makers (gdm, upward triangle, above 70% of advantageous choices in the last 20 min), intermediates (int, square, between 30S% and 70% of advantageous choices in the last 20 min), and poor decision makers (pdm, downward triangle, below 30% of advantageous choices in the last 20 min). Individual data over boxplots. **Right** Distribution density of final scores for Tph2-wt and Tph2-kd groups. **C.** Flexibility scores in the reversed-RGT corresponding to the preference for the new location of the preferred option in the RGT for gdm (upward triangle), int (square), and pdm (downward triangle). The flexibility score is the preference for the location of the non-preferred option during the RGT. Individual data over boxplots. **D. Left** Choice of the large reward option as a function of the probability of reward delivery in the probability discounting task (PDT). Lines indicate mean + SD, dashed line shows 50% chance level, Kruskal-Wallis rank sum test between all subgroups (dm-groups). **Right** Area under the curve (AUC) per individual. **E. Left** Duration of interaction in the social recognition task (SRt). Lines indicate mean + SD. Habituation with empty cage (Hab), successive encounters with same conspecific placed in the small cage (E1–3). **Right** Short-term memory ratio (STM). Boxplots classically represent the median, 25th and 75th percentiles and 1.5 IQR. *p-value < 0.05. Panels A–B: Tph2-wt, n = 60 (gdm, n = 49, int, n = 8, pdm, n = 3), Tph2-kd, n = 58 (gdm, n = 39, int, n = 10, pdm, n = 9). Panel C: Tph2-wt, n = 48 (gdm, n = 39, int, n = 6, pdm, n = 3), Tph2-kd, n = 47 (gdm, n = 31, int, n = 8, pdm, n = 8). Panel D: Tph2-wt, n = 36 (gdm, n = 30, int, n = 5, PDMs, n = 1), Tph2-kd, n = 34 (gdm, n = 22, int, n = 7, pdm, n = 5). Panel E: Tph2-wt, n = 42 (gdm, n = 34, int, n = 7, pdm, n = 1), Tph2-kd, n = 40 (gdm, n = 26, int, n = 9, pdm, n = 5).

Behavioral flexibility was similarly expressed in Tph2-wt and Tph2-kd animals (Fig.3C). Good decision makers and intermediate expressed low to high flexibility indexes whereas poor decision maker animals of both groups expressed low flexibility indexes (Fig.3C). The decreasing probability to get the large reward induced a discounting effect on the preference for the large reward (Fig.3D-left, p.anova, *probability*, F(4,256) = 153, p-value < 0.001). This discounting effect was stronger in the low-serotonin poor decision maker group than in the other groups as shown by the area under the curve (AUC) which was lower for the low-serotonin poor decision maker group (Fig. 3D-right, Kruskal-Wallis rank sum test, chi-squared = 13.02, df = 5, p-value = 0.023). All groups showed interest in the social partner (social preference, SP) indicated by the increase in interaction time from Hab to E1. Rats also formed a short-term social recognition memory (STM) of the social partner, indicated by the decrease in interaction time from E1 to E3 (Fig.3E, p.anova, *encounter*, F(3,234) = 202, p-value = 0). However, low-serotonin poor decision makers showed a lower decrease of interaction time from E1 to E3 (Fig.3E, p.anova, *encounter x treatment x RGT*, F(3,234) = 3, p-value = 0.018) and the STM ratio for this group was not different from 1 unlike the other groups (Fig.3E-right, Wilcoxon signed rank test with continuity correction, V = 9, p-value = 0.786). Odor discrimination ability was similar between Tph2-wt, Tph2-kd and subgroups (Fig.S3).

We applied a network analysis to the data to understand the relationships between the different functions of the behavioral profile of low-serotonin poor decision makers compared to other Tph2-kd and control animals. In both Tph2-kd and control groups, without poor decision makers, decision making (RGT score) and motivation for the reward (Lat i.e. latency to collect reward) were strongly connected (Fig.4A-B). However other strong pairwise connections differed between groups: in the control group a strong connection between social preference and short-term recognition memory (SP-STM) was found, in the Tph2-kd group a strong connection between short-term recognition memory and motivation for reward (STM-Lat) was found (Fig.4A-B). The low-serotonin poor decision makers (n = 5) revealed a central position of short-term social recognition (STM) with strong connections between STM and all other functions (Fig.4C).

**Figure 4.**
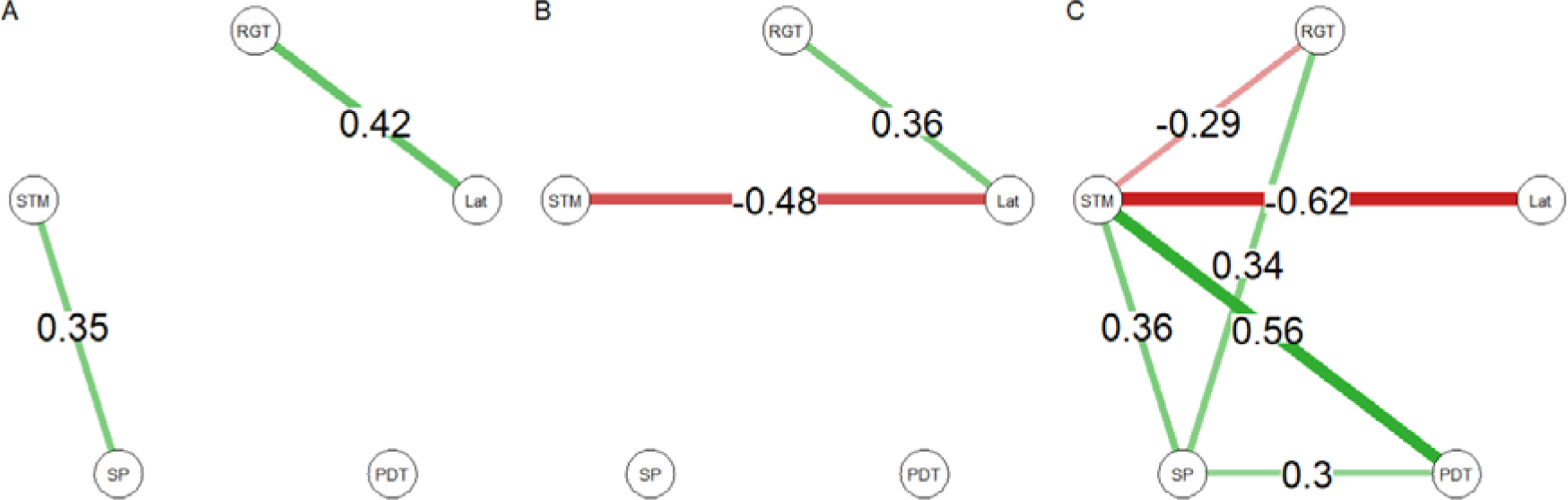
Network analysis of the behavioral profiles with Spearman’s correlations. Control (A) and Tph2-kd (B) groups without poor decision maker animals and low-serotonin poor decision makers (from the Tph2-kd) subgroup (C). Edges between cognitive functions are Spearman’s correlations, green edges for positive correlations and red edges for negative correlations. Only strong correlations (r > 0.3) are indicated and thickness of the edge the strength of the correlation. RGT: Decision making ability; Lat: motivation to collect reward; PDT: impulsive choices; SP: recognition of social novelty; STM: recognition of social familiarity. Panel A: control without poor decision makers, n = 35. Panel B: Tph2-kd without poor decision makers, n = 29. Panel C: low-serotonin poor decision makers, n = 5.

## Discussion

In this study, the acute and mild reduction of central serotonin in Teto-shTph2 rats, resulted in an increased number of individuals making disadvantageous choices in conditions of uncertainty of the RGT. Although, the link between serotonin function and poor decision making was previously shown using equivalent tests of the RGT, and systemic dietary or pharmacological approaches (de Visser et al., 2011; Koot et al., 2012). In the current study however, looking at spontaneous individual differences in decision making strategies, we show that the reduction of central serotonin levels of previously healthy individuals did not affect all animals similarly but only a subgroup of them. This difference between studies on the impact of serotonin dysfunction in all individuals (other studies) to only some individuals of the population (current study) could be due to the attention we give here to individual differences but also the advantages of using the refined model of TetO-shTph2 rat. This model offers a temporary, moderate and physiologically realistic variation of serotonin levels aimed at triggering impairments to the most spontaneously vulnerable individuals. It is also brain specific, acute (within 20 days of treatment), reversible (Sidorova et al., 2021) and it is possible to select the time window of action. This model prevent the confounding impact of developmental compensatory mechanisms of knock-out models (Alonso et al., 2023) and other off-target effects of classical non-genetic approaches.

As expected, low-serotonin poor decision makers presented a unique combination of cognitive impairments otherwise preserved in wild-type poor decision makers (Alonso et al., 2019) or in the other low-serotonin (Tph2-kd) animals (i.e. good decision makers and intermediates). Low-serotonin poor decision makers presented deficits in social recognition memory and probability-based decision making, in addition to typical hypersensitivity to reward and cognitive inflexibility which are characteristic of wild-type poor decision makers (Rivalan 2013, Alonso 2019). This combination of deficits in vulnerable individuals was specific to the acute, moderate and brain-specific decrease of serotonin function.

In the social recognition test, while low-serotonin poor decision makers presented typical social preference for novel subjects, they however maintained higher investigation time of familiar subjects, indicating a lack of habituation potentially due to a deficit in short-term memory and social recognition of familiarity. Serotonin signaling is known to control social recognition memory (Loiseau et al., 2008; Schmidt et al., 2022; Tripathy et al., 2012; Wu et al., 2021) and to be critical for the adaptation of social behaviors in home-cage environment (Alonso et al., 2023, Rivalan et al, in preparation). In degraded condition of serotonin function, vulnerable individuals might present increased difficulties in the integration and transmission of social cues to adjust behavior.

In the probability discounting task, at indifference point (P = 20), although both available options were mathematically equivalent in total amount of food they provided, low-serotonin poor decision makers, but not the other groups, switched preference for the more certain option where a (small) reward is always delivered. This behavior indicated an increased intolerance to the risk of missing a reward. Interestingly, in the RGT the poor decision makers’ hyper-sensitivity to immediate reinforcement was found to be one key driver of their choice in test (Rivalan et al., 2013). While typically, healthy poor decision makers are not impaired in the PDT and are able to choose following the absolute amount of reward of each option in this task (Alonso et al., 2023, 2019), here associated with a drop of serotonin function poor decision makers presented increased focus on short-term, immediate and certain rewards vs. long-term rewards. This is in line with the role of serotonin in anticipation of future reward and encoding of reward value (Akizawa et al., 2019; Li et al., 2016; Luo et al., 2016; Miyazaki et al., 2020).

Interestingly, despite the known effect of serotonin on behavioral flexibility (Alsiö et al., 2021; Lapiz-Bluhm et al., 2009; Wallace et al., 2014), behavioral flexibility was not worsened in the low-serotonin group compared to controls (comparing each decision making subgroups respectively). Maybe the decrease of serotonin function has to be stronger as seen in previous study and/or reversal task more complex for an effect of serotonin on behavioral flexibility to be seen. It would be interesting to test the Tph2-kd rats in other behavioral flexibility tests such as reversal learning (Izquierdo et al., 2017; Matias et al., 2017) and in more complex and ethological conditions (Nachev et al., 2021) to better understand the specific role of central serotonin in behavioral flexibility.

Here we show that a moderate, realistic and brain specific decrease of serotonin levels have a particular effect on a subset of vulnerable individuals only. In these animals, low serotonin levels affected their ability to socially habituate and to make decisions in uncertain (probabilistic) conditions by adopting a “short term” strategy focused on immediate and certain gains and in line with an increase of sensitivity to rewards. Considering the rewarding effect of social interactions (El Rawas et al., 2020), beyond impaired recognition, sustained interest for a social partner could reflect a generalization of high motivation for reward from food to social interactions. The role of serotonin in modulating social and non-social cognition is well documented (Koot et al., 2012; Mosienko et al., 2012). However, it is not yet known on which function(s) serotonin could have a primary impact and if impairments then propagate to other functions. Considering the centrality of social recognition in the behavioral network of the low-serotonin poor decision makers, an alternative view of our results could be that serotonin would primarily alter social cognition and by diffusion, the other connected cognitive functions within the network, especially risky probability-based decision making. This would not be the first time that the importance of studying social cognition as a potential origin point for the modulation of other executive and seemingly non-social functions is pointed out. In humans for instance, network approach studies have emphasized the importance of social cognition for executive function and the ability to perform daily essential activities in schizophrenia patients (Galderisi et al., 2018; Hajdúk et al., 2021). Although we did not assess decision making before inducing the decrease of serotonin, it could be assumed that a pre-existing vulnerability of the poor decision making profile could have made them more sensitive to the decrease of serotonin function and induce the impairments seen in the current study. This could explain why other low-serotonin non-poor decision makers were not similarly affected by the mild central serotonin decrease. Also following our exploratory network analysis, we propose to further investigate the properties of the behavioral network of poor decision makers in normal and “pathological” low-serotonin conditions. This would contribute to the understanding of the role of social cognition in the transition to pathological state, as a target to prevent the establishment of psychopathology. Further studies should apply the combination of the differential, multidimensional and genetic approaches presented here to model, at the individual level, transition phenomenon from adaptive to pathological states and to reveal emerging markers of pathology in spontaneously vulnerable individuals.

## Conclusion

The destabilization of the serotonergic system, using the TetO-shTph2 model, led to specific cognitive and social impairments and changes in the behavioral network of only a subset of vulnerable individuals from the general population. Besides their poor abilities to choose long-term advantageous feeding options, the low-serotonin poor decision makers expressed poor social recognition memory and strong risk aversion in a probability-based decision making task, whereas the individuals with more adapted strategy were not affected by the temporary serotonin decrease in any other tests. These results and the exploration of the cognitive network of the poor decision makers could suggest social recognition memory to be a key factor dependent on serotonin function and at the core of a larger cognitive network. With this study, we show the high potential of the Teto-shTph2 rat to model in further specific studies the transition processes of psychopathological deterioration associated with a central serotonin drop. Further longitudinal experiments in a group-housed complex semi-natural environment will increase the ethological validity of the behavioral network analysis by informing on the temporal relationships between traits specific of poor decision makers. This will also help confirm the key role of social cognition in the transition to psychopathology and challenge the role of serotonin as a triggering factor of psychopathology and contribute to the current debate on the serotonin hypothesis of psychiatric disorders (Hou et al., 2006; Moncrieff et al., 2022; Pech et al., 2018).

## Limitations of the study

We observed high variability in the levels of 5-HT, 5HIAA and TRP between batches of animals, a batch consisted of 12 animals, 6 controls and 6 Tph2-kd (Fig. S1). Descriptions of procedures we implemented to reduce variability are given in the Method section.

In this work we only studied male rats. Sex differences in decision making however exist in animals (Titulaer et al., 2012; van den Bos et al., 2012) and human with strong serotonergic neurobiological correlates (van den Bos et al., 2013). It is of prime importance to extend our study to female rats in order to evaluate the impact of serotonin decrease on the cognitive functions at the group level and on the interaction between functions, especially in female “low-serotonin poor decision makers”. It was shown recently that sex differences in decision making may be underpinned by distinct cognitive mechanisms (Truckenbrod et al., 2023) which are critical for the study of vulnerability.

## Acknowledgements

We thank Melissa Long, Susanne da Costa Goncalves, Alexej Schatz, Fatimunnisa Qadri, Niccolò Milani, Susann Matthes, Andrea Rodak, Lorenz Gygax, Vladislav Nachev for their support through technical assistance and knowledgeable discussion. We thank Annegret Dahlke, Monique Bergemann, Laura Rosenzweig, Bettina Müller, Reimunde Hellwig-Träger for their work with the animals. We thank Dalia Attalla, Alican Caglayan, Dow Glikman who made insightful comments on a previous version of the manuscript and all our colleagues of the Winter lab.

## Supplementary figures

**Figure S1.**
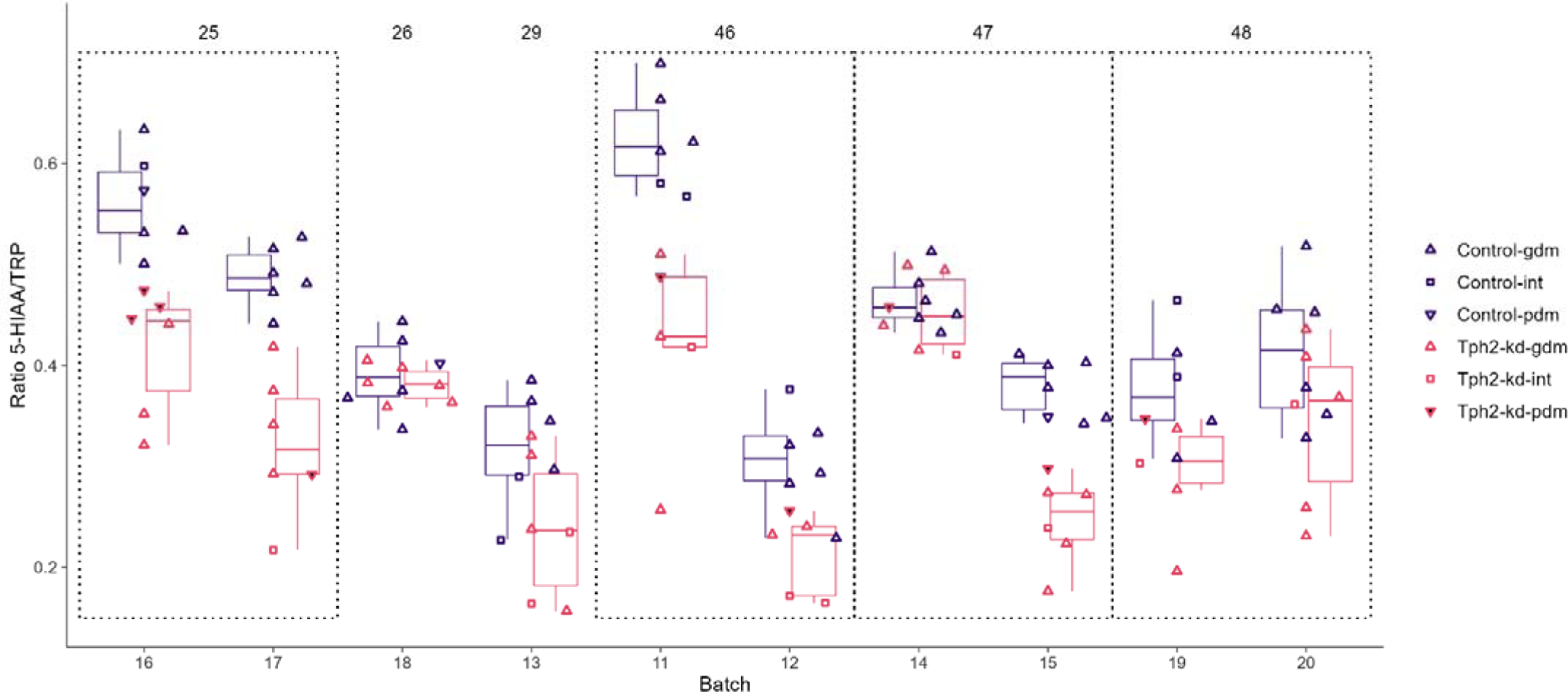
Individual ratio of 5-HIAA and tryptophan. 5-HIAA and tryptophan were more robustly detected with HPLC than 5-HT for all batches. Individual data over boxplots. The duration of treatment in days is indicated at the top of the figure and the dotted lines indicate the batches tested at the same time (pair of batches).

**Figure S2:**
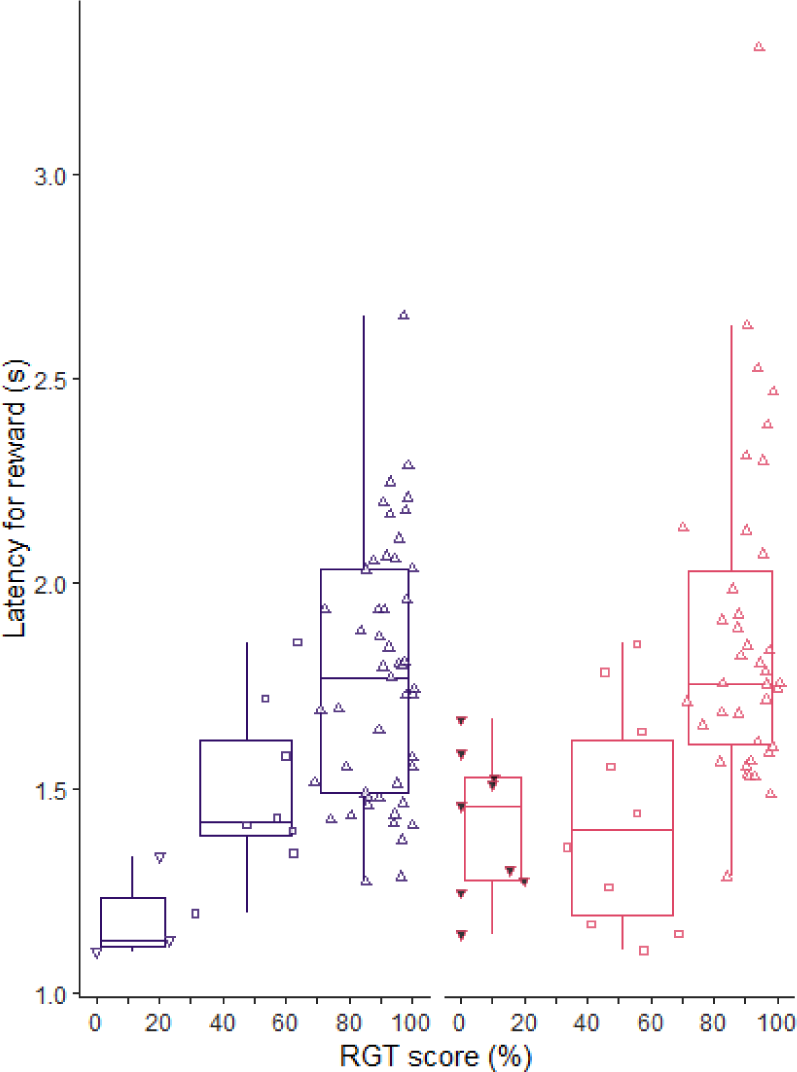
Latency to collect reward in RGT per decision making group and treatment groups. Good decision makers (GDMs, upward triangle), intermediates (INTs, square), and poor decision makers (PDMs, downward triangle). Individual data over boxplots. Tph2-wt in purple and Tph2-kd in pink. Tph2-wt, n = 60 (gdm, n = 49, int, n = 8, pdm, n = 3), Tph2-kd, n = 58 (gdm, n = 39, int, n = 10, pdm, n = 9), Tph2-kd n = 58 (gdm, n = 39, int, n = 10, pdm, n = 9), Tph2-kd, n = 58 (gdm, n = 39, int, n = 10, pdm, n = 9).

**Figure S3:**
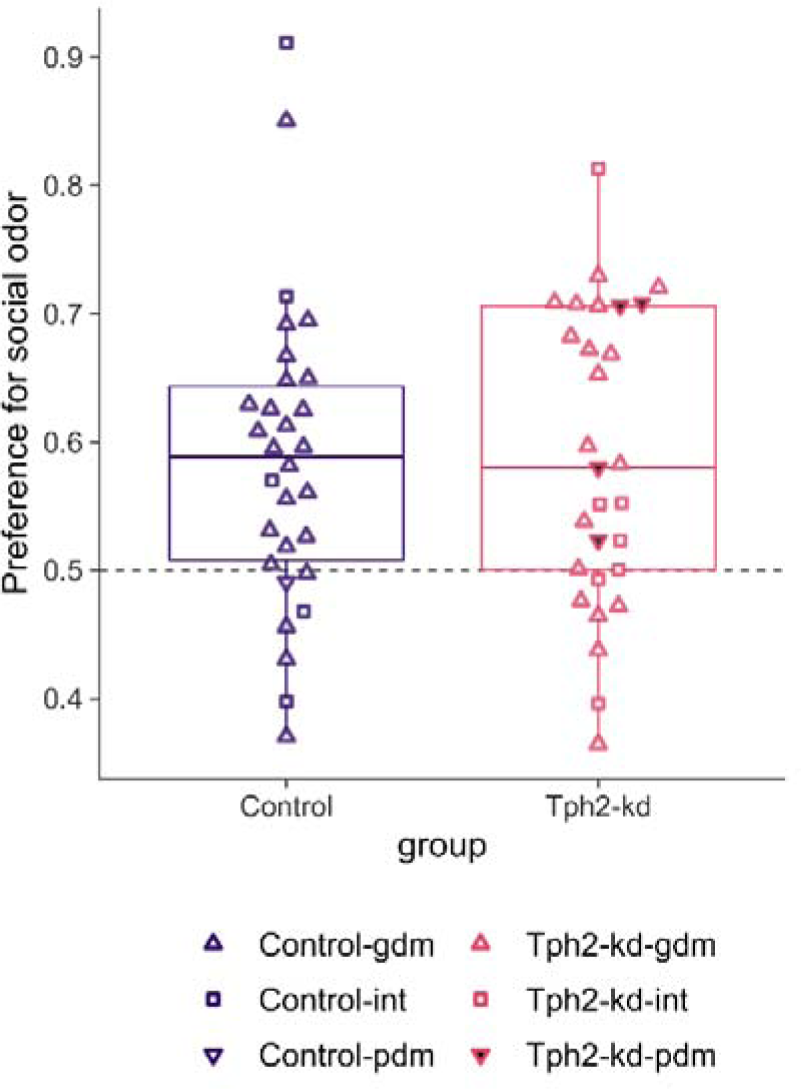
Odor preference per decision making group and treatment groups. Preference for the plate with the social odor. Good decision makers (GDMs, upward triangle), intermediates (INTs, square), and poor decision makers (PDMs, downward triangle). Individual data over boxplots. Tph2-wt in purple and Tph2-kd in pink. Tph2-wt n = 30, Tph2-kd n = 29.

## Notes

### Competing Interest Statement

The authors have declared no competing interest.

## References

Akizawa F, Mizuhiki T, Setogawa T, Takafuji M, Shidara M. 2019. The effect of 5-HT1A receptor antagonist on reward-based decision-making. J Physiol Sci JPS 69:1057–1069. doi:10.1007/s12576-019-00725-1

Alonso L, Peeva P, Ramos-Prats A, Alenina N, Winter Y, Rivalan M. 2019. Inter-individual and inter-strain differences in cognitive and social abilities of Dark Agouti and Wistar Han rats. Behav Brain Res 112188. doi:10.1016/j.bbr.2019.112188

Alonso L, Peeva P, Stasko S, Bader M, Alenina N, Winter Y, Rivalan M. 2023. Constitutive depletion of brain serotonin differentially affects rats’ social and cognitive abilities. iScience 26. doi:10.1016/j.isci.2023.105998

Alsiö J, Lehmann O, McKenzie C, Theobald DE, Searle L, Xia J, Dalley JW, Robbins TW. 2021. Serotonergic Innervations of the Orbitofrontal and Medial-prefrontal Cortices are Differentially Involved in Visual Discrimination and Reversal Learning in Rats. Cereb Cortex N Y N 1991 31:1090–1105. doi:10.1093/cercor/bhaa277

Bechara A, Damasio H. 2002. Decision-making and addiction (part I): impaired activation of somatic states in substance dependent individuals when pondering decisions with negative future consequences. Neuropsychologia 40:1675–1689. doi:10.1016/S0028-3932(02)00015-5

Blumstein DT, Daniel JC, Evans CS. 2000. JWatcher.

Borsboom D. 2017. A network theory of mental disorders. World Psychiatry Off J World Psychiatr Assoc WPA 16:5–13. doi:10.1002/wps.20375

Bui E, Nadal-Vicens M, Simon NM. 2012. Pharmacological approaches to the treatment of complicated grief: rationale and a brief review of the literature. Dialogues Clin Neurosci 14:149–157.

Cáceda R, Nemeroff CB, Harvey PD. 2014. Toward an understanding of decision making in severe mental illness. J Neuropsychiatry Clin Neurosci 26:196–213. doi:10.1176/appi.neuropsych.12110268

Dalgleish T, Black M, Johnston D, Bevan A. 2020. Transdiagnostic Approaches to Mental Health Problems: Current Status and Future Directions. J Consult Clin Psychol 88:179–195. doi:10.1037/ccp0000482

Daniel ML, Cocker PJ, Lacoste J, Mar AC, Houeto JL, Belin-Rauscent A, Belin D. 2017. The anterior insula bidirectionally modulates cost-benefit decision-making on a rodent gambling task. Eur J Neurosci 46:2620–2628. doi:10.1111/ejn.13689

de Visser L, Homberg JR, Mitsogiannis M, Zeeb FD, Rivalan M, Fitoussi A, Galhardo V, van den Bos R, Winstanley CA, Dellu-Hagedorn F. 2011. Rodent versions of the iowa gambling task: opportunities and challenges for the understanding of decision-making. Front Neurosci 5:109. doi:10.3389/fnins.2011.00109

Denburg NL, Tranel D, Bechara A. 2005. The ability to decide advantageously declines prematurely in some normal older persons. Neuropsychologia 43:1099–1106. doi:10.1016/j.neuropsychologia.2004.09.012

El Rawas R, Amaral IM, Hofer A. 2020. Social interaction reward: A resilience approach to overcome vulnerability to drugs of abuse. Eur Neuropsychopharmacol J Eur Coll Neuropsychopharmacol 37:12–28. doi:10.1016/j.euroneuro.2020.06.008

Epskamp S, Cramer AOJ, Waldorp LJ, Schmittmann VD, Borsboom D. 2012. qgraph: Network Visualizations of Relationships in Psychometric Data. J Stat Softw 48:1–18. doi:10.18637/jss.v048.i04

Fitoussi A, Le Moine C, De Deurwaerdère P, Laqui M, Rivalan M, Cador M, Dellu-Hagedorn F. 2014. Prefronto-subcortical imbalance characterizes poor decision-making: neurochemical and neural functional evidences in rats. Brain Struct Funct. doi:10.1007/s00429-014-0868-8

Galderisi S, Rucci P, Kirkpatrick B, Mucci A, Gibertoni D, Rocca P, Rossi A, Bertolino A, Strauss GP, Aguglia E, Bellomo A, Murri MB, Bucci P, Carpiniello B, Comparelli A, Cuomo A, De Berardis D, Dell’Osso L, Di Fabio F, Gelao B, Marchesi C, Monteleone P, Montemagni C, Orsenigo G, Pacitti F, Roncone R, Santonastaso P, Siracusano A, Vignapiano A, Vita A, Zeppegno P, Maj M, Italian Network for Research on Psychoses. 2018. Interplay Among Psychopathologic Variables, Personal Resources, Context-Related Factors, and Real-life Functioning in Individuals With Schizophrenia: A Network Analysis. JAMA Psychiatry 75:396–404. doi:10.1001/jamapsychiatry.2017.4607

Hajdúk M, Penn DL, Harvey PD, Pinkham AE. 2021. Social cognition, neurocognition, symptomatology, functional competences and outcomes in people with schizophrenia - A network analysis perspective. J Psychiatr Res 144:8–13. doi:10.1016/j.jpsychires.2021.09.041

Hervé M. 2020. RVAideMemoire: Testing and Plotting Procedures for Biostatistics.

Hou C, Jia F, Liu Y, Li L. 2006. CSF serotonin, 5-hydroxyindolacetic acid and neuropeptide Y levels in severe major depressive disorder. Brain Res 1095:154–158. doi:10.1016/j.brainres.2006.04.026

Hyde M, Hanson LM, Chungkham HS, Leineweber C, Westerlund H. 2015. The impact of involuntary exit from employment in later life on the risk of major depression and being prescribed anti-depressant medication. Aging Ment Health 19:381–389. doi:10.1080/13607863.2014.927821

Izquierdo A, Brigman JL, Radke AK, Rudebeck PH, Holmes A. 2017. The neural basis of reversal learning: An updated perspective. Neuroscience 345:12–26. doi:10.1016/j.neuroscience.2016.03.021

Koot S, Zoratto F, Cassano T, Colangeli R, Laviola G, van den Bos R, Adriani W. 2012. Compromised decision-making and increased gambling proneness following dietary serotonin depletion in rats. Neuropharmacology 62:1640–1650. doi:10.1016/j.neuropharm.2011.11.002

Kotnik K, Popova E, Todiras M, Mori MA, Alenina N, Seibler J, Bader M. 2009. Inducible transgenic rat model for diabetes mellitus based on shRNA-mediated gene knockdown. PloS One 4:e5124. doi:10.1371/journal.pone.0005124

Lapiz-Bluhm MDS, Soto-Piña AE, Hensler JG, Morilak DA. 2009. Chronic intermittent cold stress and serotonin depletion induce deficits of reversal learning in an attentional set-shifting test in rats. Psychopharmacology (Berl) 202:329–341. doi:10.1007/s00213-008-1224-6

Li Y, Zhong W, Wang D, Feng Q, Liu Z, Zhou J, Jia C, Hu F, Zeng J, Guo Q, Fu L, Luo M. 2016. Serotonin neurons in the dorsal raphe nucleus encode reward signals. Nat Commun 7:10503. doi:10.1038/ncomms10503

Loiseau F, Dekeyne A, Millan MJ. 2008. Pro-cognitive effects of 5-HT6 receptor antagonists in the social recognition procedure in rats: implication of the frontal cortex. Psychopharmacology (Berl) 196:93–104. doi:10.1007/s00213-007-0934-5

Luo M, Li Y, Zhong W. 2016. Do dorsal raphe 5-HT neurons encode “beneficialness”? Neurobiol Learn Mem 135:40–49. doi:10.1016/j.nlm.2016.08.008

Matias S, Lottem E, Dugué GP, Mainen ZF. 2017. Activity patterns of serotonin neurons underlying cognitive flexibility. eLife 6. doi:10.7554/eLife.20552

Matthes S, Mosienko V, Popova E, Rivalan M, Bader M, Alenina N. 2019. Targeted Manipulation of Brain Serotonin: RNAi-Mediated Knockdown of Tryptophan Hydroxylase 2 in Rats. ACS Chem Neurosci. doi:10.1021/acschemneuro.8b00635

Miyazaki K, Miyazaki KW, Sivori G, Yamanaka A, Tanaka KF, Doya K. 2020. Serotonergic projections to the orbitofrontal and medial prefrontal cortices differentially modulate waiting for future rewards. Sci Adv 6:eabc7246. doi:10.1126/sciadv.abc7246

Moncrieff J, Cooper RE, Stockmann T, Amendola S, Hengartner MP, Horowitz MA. 2022. The serotonin theory of depression: a systematic umbrella review of the evidence. Mol Psychiatry 1–14. doi:10.1038/s41380-022-01661-0

Moriana JA, Gálvez-Lara M, Corpas J. 2017. Psychological treatments for mental disorders in adults: A review of the evidence of leading international organizations. Clin Psychol Rev 54:29–43. doi:10.1016/j.cpr.2017.03.008

Mosienko V, Bert B, Beis D, Matthes S, Fink H, Bader M, Alenina N. 2012. Exaggerated aggression and decreased anxiety in mice deficient in brain serotonin. Transl Psychiatry 2:e122. doi:10.1038/tp.2012.44

Mu L, Wang J, Cao B, Jelfs B, Chan RHM, Xu X, Hasan M, Zhang X, Li Y. 2015. Impairment of cognitive function by chemotherapy: association with the disruption of phase-locking and synchronization in anterior cingulate cortex. Mol Brain 8:32. doi:10.1186/s13041-015-0125-y

Nachev V, Rivalan M, Winter Y. 2021. Two-dimensional reward evaluation in mice. Anim Cogn 24:981–998. doi:10.1007/s10071-021-01482-8

Nussbaumer-Streit B, Thaler K, Chapman A, Probst T, Winkler D, Sönnichsen A, Gaynes BN, Gartlehner G. 2021. Second-generation antidepressants for treatment of seasonal affective disorder. Cochrane Database Syst Rev 3:CD008591. doi:10.1002/14651858.CD008591.pub3

Opsahl T, Agneessens F, Skvoretz J. 2010. Node centrality in weighted networks: Generalizing degree and shortest paths. Soc Netw 32:245–251. doi:10.1016/j.socnet.2010.03.006

Pech J, Forman J, Kessing LV, Knorr U. 2018. Poor evidence for putative abnormalities in cerebrospinal fluid neurotransmitters in patients with depression versus healthy non-psychiatric individuals: A systematic review and meta-analyses of 23 studies. J Affect Disord 240:6–16. doi:10.1016/j.jad.2018.07.031

Pittaras E, Callebert J, Chennaoui M, Rabat A, Granon S. 2016. Individual behavioral and neurochemical markers of unadapted decision-making processes in healthy inbred mice. Brain Struct Funct 221:4615–4629. doi:10.1007/s00429-016-1192-2

Proctor D, Williamson RA, Latzman RD, de Waal FBM, Brosnan SF. 2014. Gambling primates: reactions to a modified Iowa Gambling Task in humans, chimpanzees and capuchin monkeys. Anim Cogn 17:983–995. doi:10.1007/s10071-014-0730-7

R Core Team. 2019. R: The R Project for Statistical Computing, R Foundation for Statistical Computing, Vienna, Austria. https://www.R-project.org/. https://www.r-project.org/

Rivalan M, Ahmed SH, Dellu-Hagedorn F. 2009. Risk-prone individuals prefer the wrong options on a rat version of the Iowa Gambling Task. Biol Psychiatry 66:743–749. doi:10.1016/j.biopsych.2009.04.008

Rivalan M, Coutureau E, Fitoussi A, Dellu-Hagedorn F. 2011. Inter-individual decision-making differences in the effects of cingulate, orbitofrontal, and prelimbic cortex lesions in a rat gambling task. Front Behav Neurosci 5:22. doi:10.3389/fnbeh.2011.00022

Rivalan M, Valton V, Seriès P, Marchand AR, Dellu-Hagedorn F. 2013. Elucidating Poor Decision-Making in a Rat Gambling Task. PLoS ONE 8:e82052. doi:10.1371/journal.pone.0082052

Schmidt SD, Zinn CG, Cavalcante LE, Ferreira FF, Furini CRG, Izquierdo I, de Carvalho Myskiw J. 2022. Participation of Hippocampal 5-HT5A, 5-HT6 and 5-HT7 Serotonin Receptors on the Consolidation of Social Recognition Memory. Neuroscience S0306-4522(22)00308–6. doi:10.1016/j.neuroscience.2022.06.016

Sidorova M, Kronenberg G, Matthes S, Petermann M, Hellweg R, Tuchina O, Bader M, Alenina N, Klempin F. 2021. Enduring Effects of Conditional Brain Serotonin Knockdown, Followed by Recovery, on Adult Rat Neurogenesis and Behavior. Cells 10:3240. doi:10.3390/cells10113240

Steingroever H, Wetzels R, Horstmann A, Neumann J, Wagenmakers E-J. 2013. Performance of healthy participants on the Iowa Gambling Task. Psychol Assess 25:180–193. doi:10.1037/a0029929

Titulaer M, van Oers K, Naguib M. 2012. Personality affects learning performance in difficult tasks in a sex-dependent way. Anim Behav 83:723–730. doi:10.1016/j.anbehav.2011.12.020

Tripathy R, McHugh RJ, Bacon ER, Salvino JM, Morton GC, Aimone LD, Huang Z, Mathiasen JR, DiCamillo A, Huffman MJ, McKenna BA, Kopec K, Lu LD, Qian J, Angeles TS, Connors T, Spais C, Holskin B, Duzic E, Schaffhauser H, Rossé GC. 2012. Discovery of 7-arylsulfonyl-1,2,3,4, 4a,9a-hexahydro-benzo[4,5]furo[2,3-c]pyridines: identification of a potent and selective 5-HT₆ receptor antagonist showing activity in rat social recognition test. Bioorg Med Chem Lett 22:1421–1426. doi:10.1016/j.bmcl.2011.12.026

Truckenbrod LM, Cooper EM, Orsini CA. 2023. Cognitive mechanisms underlying decision making involving risk of explicit punishment in male and female rats. Cogn Affect Behav Neurosci 23:248–275. doi:10.3758/s13415-022-01052-6

van den Bos R, Homberg J, de Visser L. 2013. A critical review of sex differences in decision-making tasks: focus on the Iowa Gambling Task. Behav Brain Res 238:95–108. doi:10.1016/j.bbr.2012.10.002

van den Bos R, Jolles J, van der Knaap L, Baars A, de Visser L. 2012. Male and female Wistar rats differ in decision-making performance in a rodent version of the Iowa Gambling Task. Behav Brain Res 234:375–379. doi:10.1016/j.bbr.2012.07.015

Wallace A, Pehrson AL, Sánchez C, Morilak DA. 2014. Vortioxetine restores reversal learning impaired by 5-HT depletion or chronic intermittent cold stress in rats. Int J Neuropsychopharmacol 17:1695–1706. doi:10.1017/S1461145714000571

Wheeler B, Torchiano M. 2016. lmPerm: Permutation Tests for Linear Models.

World Health Organization MH and SU. 2021. Comprehensive Mental Health Action Plan 2013-2030.

Wu X, Morishita W, Beier KT, Heifets BD, Malenka RC. 2021. 5-HT modulation of a medial septal circuit tunes social memory stability. Nature 599:96–101. doi:10.1038/s41586-021-03956-8

Zoratto F, Sinclair E, Manciocco A, Vitale A, Laviola G, Adriani W. 2014. Individual differences in gambling proneness among rats and common marmosets: an automated choice task. BioMed Res Int 2014:927685. doi:10.1155/2014/927685

